# Relatedness and genetic structure of big brown bat (*Eptesicus fuscus*) maternity colonies in an urban-wildland interface provides insight into rabies transmission dynamics

**DOI:** 10.1101/2020.06.01.128496

**Authors:** Faith M. Walker, Colin J. Sobek, Camille E. Platts-McPharlin, Carol L. Chambers

## Abstract

Big brown bats (*Eptesicus fuscus*) are the bat species most frequently found to be rabid in North America and are a key source of sylvatic rabies in wildlife. Females can form summer maternity colonies in urban areas, often using access holes in the exterior of buildings to roost in relatively large numbers. In Flagstaff, Arizona, these roosts are commonly found in houses adjacent to golf courses, where habitat quality (food, water, shelter) is high for bats and for mesocarnivores such as striped skunks (*Mephitis mephitis*) and gray foxes (*Urocyon cinereoargenteus*). Periodic rabies outbreaks in Flagstaff involving all three of these mammals are primarily caused by an *E. fuscus* variant of the disease. However, little is known about *E. fuscus* social behavior during the summer months and how it may drive space use and hence disease exposure to conspecifics and mesocarnivores. To address this knowledge gap, we collected 88 unique genetic samples via buccal swabs from *E. fuscus* captured at four maternity roosts surrounding a golf course during summer of 2013. We used 7 microsatellite loci to estimate genetic relatedness among individuals and genetic structure within and among colonies in order to infer whether females selected roosts based on kinship, and used genetics and radio telemetry to determine the frequency of roost switching. We found roost switching through genetics (two mother and adult daughter pairs at the same and different roosts) and telemetry, and no evidence of elevated genetic relatedness within colonies or genetic structure between colonies. These results have important implications for disease transmission dynamics in that social cohesion based on relatedness does not act to constrain the virus to a particular roost area. Instead, geographic mobility may increase disease exposure to neighboring areas. We discuss mitigating actions for bat conservation and public health.

## Introduction

Of infectious diseases, rabies virus (RABV) has the highest fatality rate, resulting in nearly 60,000 human deaths worldwide each year (WHO 2018). Dogs are the predominant RABV vector globally, but in the Americas, where vaccination programs target domestic carnivores, bat-mediated rabies is responsible for the majority of human rabies cases (Vigilato et al. 2013). In North America, big brown bats (*Eptesicus fuscus*) are the species associated with the most human exposures because they often roost in houses and other structures, and because of their large geographic distribution (Agosta 2002). Although transmission of RABV from *E. fuscus* to humans only occurs rarely, thanks at least in part to post-exposure prophylaxis, transmission to wildlife species such as carnivores occur more frequently (Kuzmin et al. 2012). Arizona consistently ranks among the prominent states for RABV positive wildlife, with bats and skunks being the most common reservoirs (Ma et al. 2018, ADHS 2020).

The urban-wildland interface presents enhanced opportunities for sylvatic RABV transmission. Flagstaff, Arizona, is a high elevation city (2170 m) of 75,000 people and is surrounded by 1.86 million acres of national forest. It has been experiencing periodic RABV outbreaks in bats, striped skunks (*Mephitis mephitis*) and gray foxes (*Urocyon cinereoargenteus*) for at least 15 years. Most of these outbreaks have been driven by the *E. fuscus* RABV variant introduced into the two mesocarnivore species’ populations, independently, during each outbreak (Kuzmin et al. 2012). Although the route of transmission between bats, skunks, and foxes is unknown, it is hypothesized that suburban wildlife/human interface zones, such as golf courses, provide habitat for these species due to abundant water (ponds), food (bats: insects; mesocarnivores: bird feeders, pet food), and shelter (infrequently used homes), hence providing opportunities for spillover from bat bites or ingestion of infected bat carcasses (Theimer et al. 2015, Theimer et al. 2017a, Theimer et al. 2017b). In summer, female *E. fuscus* form maternity roosts in houses neighboring golf courses, many of which are second homes and hence not used frequently by humans. Bats roost and raise young in siding and other external entrances, and these neighborhoods with access to rich resources become potential enzootic rabies hotspots. As the weather cools in October, bats hibernate or depart, possibly to lower elevation, until the following May.

Relatedness within urban *E. fuscus* maternity roosts and the amount of roost switching during summer months will provide clues about intra-species transmission dynamics of rabies, which may assist with management to mitigate and model rabies outbreaks. It is unknown whether maternity roosts within buildings are composed of groups of related females, or whether they have high fidelity to roosts while raising offspring or commonly switch roosts. Fidelity to roosts or high relatedness within a colony would suggest that rabies outbreaks could be limited to the roost area rather than spread throughout a larger area involving multiple roosts. Brigham and Fenton (2011) found that female *E. fuscus* that inhabited maternity roosts in buildings in Ottawa, Canada, exhibited high site fidelity and suggested that the roosts may be composed of cohesive social groups rather than random assortment of individuals. In forest-dwelling *E. fuscus* in Saskatchewan, Canada, Willis and Brigham (2004) found that females were loyal to a small area within years, and that in some cases this extended between years. They also found non-random roosting associations, with 40% of bats roosting with the same associates the following year and pairs often switching roosts together. Here, we assessed roost fidelity and kinship in the Flagstaff golf course maternity colony system, hypothesizing based on the above studies that relatedness is higher within than among colonies and roost-switching is infrequent. We evaluated short-term movements by tracking individuals with microsatellite DNA and radio telemetry, and determined whether longer-term kin relationships were present by evaluating parentage, genetic relatedness, and genetic structure within and among colonies.

## Methods

### Sampling and radio telemetry

We captured female *E. fuscus* from four maternity roosts in the fascia of houses and nearby ponds in a 1.7 km^2^ area of Continental Country Club area of Flagstaff, Arizona (35°11′57″N 111°37′52″W), between May and August 2012 and 2013. Telemetry data were collected in 2012, and guided roost selection for genetic data collection in 2013. For houses, nets with funnels (Figure 1) were deployed at dusk and left open for up to four hours, and all or nearly all (>90%) bats at roosts were captured. Each roost was netted once. At ponds, we captured bats using mist nets. Upon bat capture, we collected a buccal swab (Whatman Omniswabs, Whatman International Ltd., Maidstone, UK) for genetic analysis. We gently rotated swabs in the mouth for 1 minute before placing them in 1.5 mL tubes containing 500 μL of RNAlater (Ambion, Austin, TX). Total time of bat capture was 30 minutes or less; samples were stored at −80°C until DNA extraction. We captured and handled bats under guidelines of the American Society of Mammalogists (Sikes 2016) and with approval of Northern Arizona University’s Institutional Animal Care and Use Committee (protocol numbers 07-006-R1, 07-006-R2). No bat suffered injury or mortality as part of this study.

**Figure 1.**
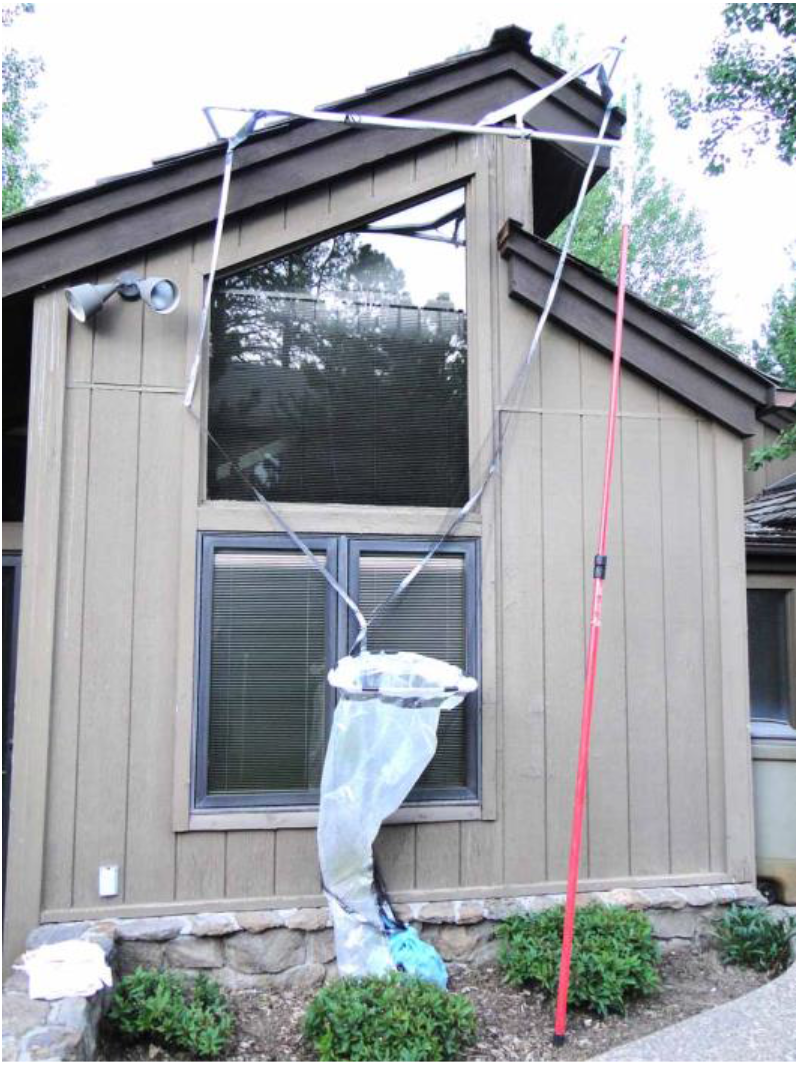
Net apparatus used to capture *Eptesicus fuscus* maternity colonies roosting in the fascia of houses. The apparatus consisted of netting supported by PVC piping, and a shower curtain acting as a funnel from the net to a bat handler. Prior to netting, exits were observed to determine access point for bats.

We attached a radio transmitter (BD-2 model, Holohil Systems Ltd., Carp, Ontario, Canada) using non-toxic latex glue between the scapulae of two female *E. fuscus* captured at a pond. All transmitters weighed ≤5% of the mass of the bat (Neubaum et al. 2005). We located each bat using a handheld directional H antenna (RA-23K, Telonics, Mesa, Arizona, USA) or omnidirectional car-top whip antenna (RA-5A, Telonics, Mesa, Arizona, USA) until transmitters fell off (7 to 14 days), identifying roosts when we found the radio-tagged animal.

### Genotyping and locus behavior

We extracted genomic DNA following the buccal swab protocol described in Walker et al. (2016). Seven microsatellite loci (MMG9, (Castella and Ruedi 2000); COTOF09, (Piaggio et al. 2009); EUMA18, EUMA29, EUMA39, EUMA43, EUMA55, (Walker et al. 2014)) were PCR-amplified on MJ Research PTC-200 thermal cyclers in 15 μL reactions. Reactions contained 1X Mg-free PCR buffer (Invitrogen), 2 mM MgCl_2_ (Invitrogen), 0.2 mM dNTPs, 0.1 U/μL Platinum Taq DNA polymerase (Invitrogen), 0.04 μg/μL Ultrapure non-acetylated Bovine Serum Albumin (Ambion), 0.2 μM fluorescently-labelled forward primer, 0.2 μM reverse primer, H_2_O, and 5 μL DNA template. We used a standard annealing temperature of 54°C for all loci except EUMA18, EUMA39, and EUMA43, which had an annealing temperature of 61°C (Walker et al. 2014). Cycling conditions began with a denaturation step of 94°C for 2 min, followed by 35 cycles of 30 s at 94°C, 30 s at 54 or 61°C, and 1 min at 72°C, then concluded with a final extension step of 72°C for 10 min. PCR products were diluted 1:50 prior to fragment analysis on an Applied Biosystems 3130 Genetic Analyzer. We used GeneMapper 4.0 software to score alleles, and GenAlEx 6.5 (Peakall and Smouse 2012) to identify recaptured bats.

We examined deviations from Hardy–Weinberg equilibrium and linkage disequilibrium between pairs of loci with program GENEPOP 4.7.3 (Rousset 2008). In addition to the non-directional Hardy-Weinberg test, we assessed heterozygote or homozygote deficits, and we applied a sequential Bonferroni procedure (Rice 1989) to adjust for multiple tests where appropriate.

### Genetic relatedness and structure within and between roosts

We employed genotypic data to calculate genetic relatedness (Queller and Goodnight 1989) between pairs of individuals in each roost using GenAlEx 6.5 (Peakall and Smouse 2012). Randomization tests were performed to determine whether there was a difference in pairwise relatedness between the three largest roosts (10,000 permutations via Resampling Stats 4.0; (Simon 1990). We investigated population structure between the three largest roosts with analysis of molecular variance (AMOVA) via GenAlEx 6.5 (Peakall and Smouse 2012), which uses permutation methods (999 iterations) to test for significance of the variance components of within and among-roost levels of genetic structure. We also assessed genetic differentiation using FSTAT 2.9.3 (Goudet 2001) to calculate Nei’s genetic distance between roost pairs and to examine significance via 1,000 permutations of the data (P = 0.017, adjusted for multiple comparisons).

### Parentage estimation

We used the log-likelihood-based program Cervus 3.0 (Kalinowski et al. 2007) to identify mother-adult daughter pairs. The average probability of excluding a randomly-selected unrelated bat from parentage was > 0.999. Because we had no prior knowledge of maternity, each female was both a candidate offspring and a candidate mother, in 100,000 simulations. Parameters included 45% of candidate mothers sampled, 100% of loci genotyped, and an error rate of 2%. Parentage was assigned when all of the following criteria were satisfied: 1) LOD > 3.0 (Marshall et al. 1998); 2) delta > 95%; and, 3) no allelic mismatches.

## Results

### Captures at roosts

We captured and collected genetic samples from 92 female *E. fuscus* from four roosts and nearby ponds, representing 88 unique individuals and four recaptures. Three recaptures were at different roosts, and one was at a pond. The number of bats at each roost were 28, 22, 18, and 9, with an additional 11 at ponds. Females were in various reproductive conditions of gestating (N= 38), lactating (N=30), and non-reproductive (N=14); 6 were unknown.

### Locus behavior

All loci adhered to Hardy-Weinberg expectations for the probability test as well as for heterozygote or homozygote deficits, except for loci EUMA39 and EUMA43, which were dropped from further analyses. No locus pairs were in linkage disequilibrium, and there was no evidence for null alleles for any locus. Mean expected heterozygosity was 0.835 (± 0.19) and the mean number of alleles per locus was 11. Multilocus genotypes are available in Supplementary Table 1.

### Genetic relatedness and structure

Mean pairwise genetic relatedness was no higher within than among maternity colonies (*P* > 0.12). We found no significant genetic structure among the three largest maternity colonies (AMOVA, Table 1; Nei’s genetic distance, Table 2).

**Table 1.** Within and among-roost levels of genetic variance, as indicated by Analysis of Molecular Variance (AMOVA) for the three largest *E. fuscus* maternity roosts. Ninety-nine percent of genetic variance was within roosts, with only 1% among roosts.

| Source | df | SS | MS | Est. Var. | % | Statistic | P-value |
| --- | --- | --- | --- | --- | --- | --- | --- |
| Among Pops | 2 | 10.658 | 5.329 | 0.039 | 1% | PhiPT | 0.158 |
| Within Pops | 65 | 289.342 | 4.451 | 4.451 | 99% |  |  |
| Total | 67 | 300.000 |  | 4.491 | 100% |  |  |

**Table 2.**
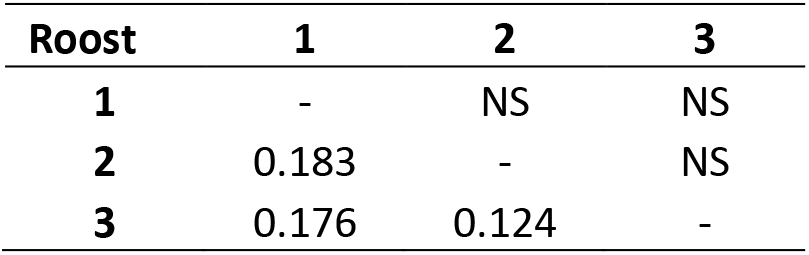
Nei’s genetic distance (below diagonal) and statistical significance (above diagonal) between pairs of *E. fuscus* roosts in the Country Club area of Flagstaff, Arizona, in 2013.

### Analysis of parentage

The delta criterion was 5.72 at the 95% confidence level and 3.62 at the 80% confidence level. Two mother-daughter pairs met the mother-daughter criteria at the 95% confidence level: one pair was captured at the same roost, and members of the other pair were captured at different roosts. Of the 6 pairs at the 80% confidence level, 67% were from different roosts. All members of pairs were adults.

### Movement between roosts

In addition to the adult mother-daughter pair detected at different roosts, we found via genetic recaptures that three females in varying reproductive condition moved between multiple roosts that were 0.27 and 0.12 km apart (Figure 2). Using telemetry, we also identified two cases of females switching roosts that were 153 m and 523 m apart. A pregnant bat used a roost for two days before shifting to a second roost for two days and a non-reproductive female used the same roost for four days before shifting to a second roost. Three roosts were in houses (under eaves, within chimney caps); one roost was an artificial roost attached to the side of a house.

**Figure 2.**
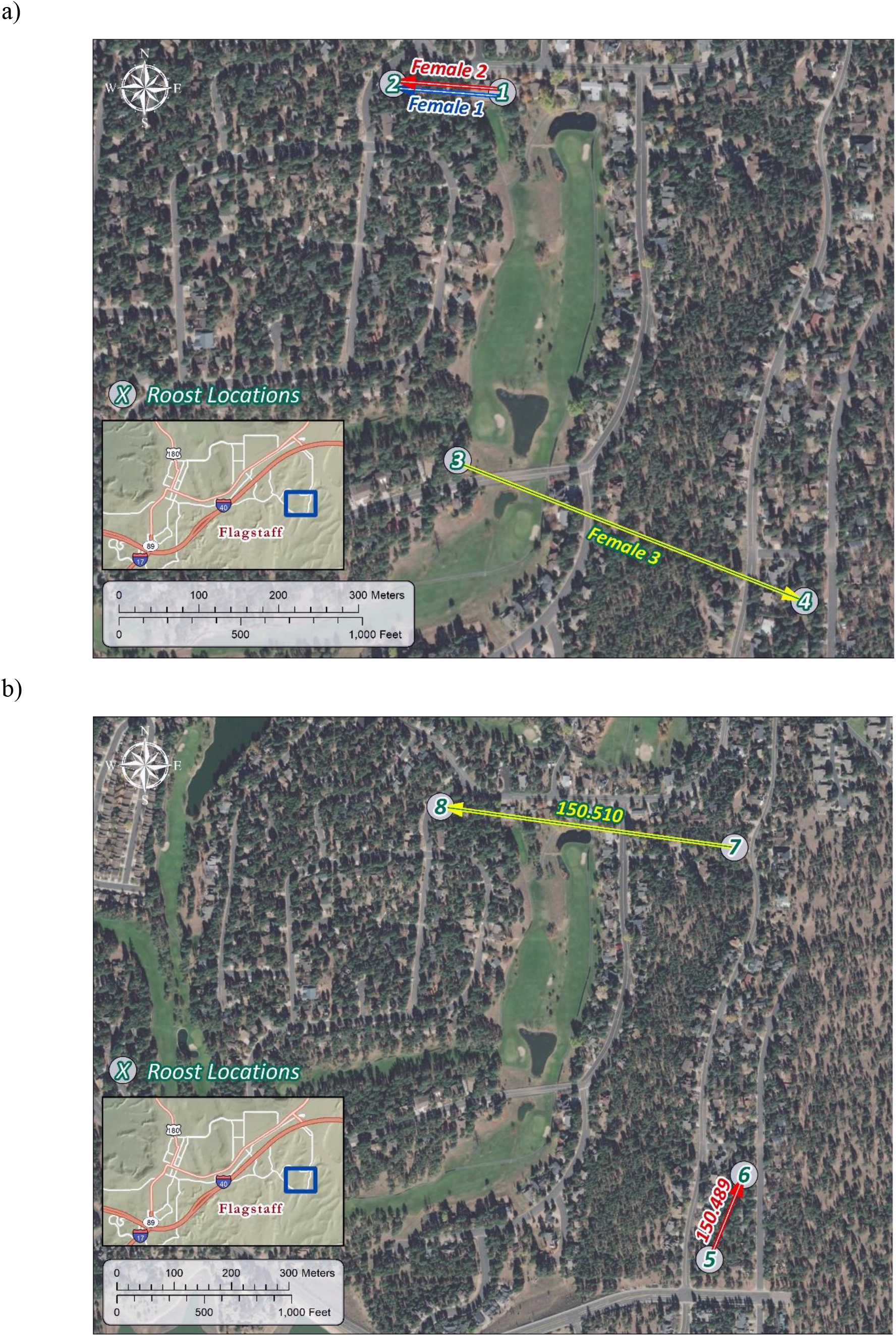
Roost switches by female *Eptesicus fuscus* in Flagstaff, Arizona, as determined by a) genetics, and b) radio-telemetry. Background imaging source: Esri, DigitalGlobe, GeoEye, Earthstar Geographics, CNES/Airbus DS, USDA, USGS, AeroGRID, IGN, and the GIS User Community.

## Discussion

The lack of elevated genetic relatedness within *E. fuscus* maternity colonies, lack of genetic structure among colonies, detection of mothers and their adult daughters at both the same and different maternity roosts, and observations of roost switching all suggest a high level of movement. Females at this location are not selecting their summer maternity roosts based on kinship. Other factors can drive roosting decisions in this species (e.g., ambient temperature, Ellison et al. 2007), and other Chiroptera (Lewis 1995, Fagan et al. 2018). That said, the return of a mother and adult daughter to the same summer maternity colony may indicate that active preferential associations can be present (Willis and Brigham 2004), or it may be a passive association generated by attraction to the same roost.

Our findings of interactions between colonies and the geographic mobility of females at the time of year that they occupy a location in Flagstaff’s urban-wildland interface that is also attractive to mesocarnivores implies an increased risk of sylvatic spread of RABV in a larger geographic area than a single roost. Webber et al. (2016) used social network analyses to predict hypothetical pathogen transmission in *E. fuscus* maternity colonies. Their models indicated that a pathogen could move significantly faster in a building roost than a tree roost. However, the common practice of excluding bats from building roosts may not be advantageous from a public health perspective, since studies have found that *E. fuscus* increased movements after roost eviction (Brigham and Fenton 2011, Streicker et al. 2013), with bats settling in new roosts within 1 km (Brigham and Fenton 2011). Increased intermixing and movements caused pathogen spread into neighboring areas (Donnelly et al. 2006, Woodroffe et al. 2006). A larger telemetry study to also tracks bats in areas of the country club adjoining our <2km^2^ site would determine the geographic extent of movements across the area and the scale of fidelity to particular areas (Willis and Brigham 2004).

Exclusion of *E. fuscus* before pups were born was associated with a decrease in reproduction (Brigham and Fenton 2011). Given the current stresses on North American bats (e.g., white-nose syndrome, (Frick et al. 2010)), it is important to align public health and conservation priorities. Streicker et al. (2013) suggested that installation of artificial roosts in urban areas would decrease risks to humans and companion animals. Further, they found that bat boxes may reduce roost switching after exclusion from buildings, thereby minimizing RABV transmission to neighboring colonies and to mesocarnivores in these areas. Eliminating den sites of skunks via sealing access under houses, decreasing pet food availability, and baiting with oral rabies vaccines under bird feeders are potential measures for reducing bat-mesocarnivore transmission (Theimer et al. 2015, Theimer et al. 2017a, Theimer et al. 2017b). These measures, in addition to maintaining current rabies vaccinations of pets, will assist with mitigating the implications of this study: summer maternity colonies of *E. fuscus* are vagile, with the potential to circulate RABV to neighboring areas.

## Acknowledgments

Funding was provided by the the State of Arizona Technology and Research Initiative Fund, Arizona Biomedical Research Commission, USDA APHIS Wildlife Services, and a Northern Arizona University Hooper Undergraduate Research Award (to CEP). Thanks to E. Saunders-Considine and B., A., R., and T. Keeley for assistance with mist netting and radio tracking, and J. Jenness for map creation.

## References cited

ADHS. 2020. Arizona Department of Health Services: Rabies data, publications, and maps. https://www.azdhs.gov/preparedness/epidemiology-disease-control/rabies/#data-publications-maps.

Agosta, S. J. 2002. Habitat use, diet and roost selection by the big brown bat (*Eptesicus fuscus*) in North America: a case for conserving an abundant species. Mammal Review 32:179–198.

Brigham, R., and B. Fenton. 2011. The influence of roost closure on the roosting and foraging behaviour of *Eptesicus fuscus* (Chiroptera: Vespertilionidae). Canadian Journal of Zoology 64:1128–1133.

Castella, V., and M. Ruedi. 2000. Characterization of highly variable microsatellite loci in the bat *Myotis myotis* (Chiroptera: Vespertilionidae). Molecular Ecology 9:1000–1002.

Donnelly, C. A., R. Woodroffe, D. R. Cox, F. J. Bourne, C. L. Cheeseman, R. S. Clifton-Hadley, G. Wei, G. Gettinby, P. Gilks, H. Jenkins, W. T. Johnston, A. M. Le Fevre, J. P. McInerney, and W. I. Morrison. 2006. Positive and negative effects of widespread badger culling on tuberculosis in cattle. Nature 439:843–846.

Fagan, K. E., E. V. Willcox, L. T. Tran, R. F. Bernard, and W. H. Stiver. 2018. Roost selection by bats in buildings, Great Smoky Mountains National Park. The Journal of Wildlife Management 82:424–434.

Frick, W. F., J. F. Pollock, A. C. Hicks, K. E. Langwig, D. S. Reynolds, G. G. Turner, C. M. Butchkoski, and T. H. Kunz. 2010. An emerging disease causes regional population collapse of a common North American bat species. Science 329:679–682.

Goudet, J. 2001. FSTAT, a program to estimate and test gene diversities and fixation indices (version 293).

Kalinowski, S. T., M. L. Taper, and T. C. Marshall. 2007. Revising how the computer program cervus accommodates genotyping error increases success in paternity assignment. Molecular Ecology 16:1099–1106.

Kuzmin, I. V., M. Shi, L. A. Orciari, P. A. Yager, A. Velasco-Villa, N. A. Kuzmina, D. G. Streicker, D. L. Bergman, and C. E. Rupprecht. 2012. Molecular inferences suggest multiple host shifts of rabies viruses from bats to mesocarnivores in Arizona during 2001–2009. PLOS Pathogens 8:e1002786.

Lewis, S. E. 1995. Roost fidelity of bats: A review. Journal of Mammalogy 76:481–496.

Ma, X., B. P. Monroe, J. M. Cleaton, L. A. Orciari, P. Yager, Y. Li, J. D. Kirby, J. D. Blanton, B. W. Petersen, and R. M. Wallace. 2018. Rabies surveillance in the United States during 2016. J Am Vet Med Assoc 252:945–957.

Neubaum, D. J., M. A. Neubaum, L. E. Ellison, and T. J. O’Shea. 2005. Survival and condition of big brown bats (*Eptesicus fuscus*) after radiotagging. Journal of Mammalogy 86:95–98.

Peakall, R., and P. E. Smouse. 2012. GenAlEx 6.5: genetic analysis in Excel. Population genetic software for teaching and research--an update. Bioinformatics (Oxford, England) 28:2537–2539.

Piaggio, A. J., K. E. G. Miller, M. D. Matocq, and S. L. Perkins. 2009. Eight polymorphic microsatellite loci developed and characterized from Townsend’s big-eared bat, *Corynorhinus townsendii*. Molecular Ecology Resources 9:258–260.

Queller, D., and K. Goodnight. 1989. Estimating relatedness using genetic markers. Evolution 43:258–275.

Rice, W. R. 1989. Analyzing tables of statistical tests. Evolution 43:223–225.

Rousset, F. 2008. genepop’007: a complete re-implementation of the genepop software for Windows and Linux. Molecular Ecology Resources 8:103–106.

Sikes, R. S. 2016. Guidelines of the American Society of Mammalogists for the use of wild mammals in research and education. Journal of Mammalogy 97:663–688.

Simon, J. 1990. Resampling Stats 4.0. in I. Resampling Stats, editor., http://www.resample.com.

Streicker, D. G., R. Franka, F. R. Jackson, and C. E. Rupprecht. 2013. Anthropogenic roost switching and rabies virus dynamics in house-roosting big brown bats. Vector Borne Zoonotic Dis 13:498–504.

Theimer, T., T. Talk, S. Johnson, and D. L. Bergman. 2017a. Bird feeders as locations for skunk uptake of oral rabies vaccine baits. Journal of Wildlife Diseases 53:424–427.

Theimer, T. C., A. C. Clayton, A. Martinez, D. L. Peterson, and D. L. Bergman. 2015. Visitation rate and behavior of urban mesocarnivores differs in the presence of two common anthropogenic food sources. Urban Ecosystems 18:895–906.

Theimer, T. C., A. C. Dyer, B. W. Keeley, A. T. Gilbert, and D. L. Bergman. 2017b. Ecological potential for rabies virus transmission via scavenging of dead bats by mesocarnivores. J Wildl Dis 53:382–385.

Vigilato, M. A. N., O. Cosivi, T. Knöbl, A. Clavijo, and H. M. T. Silva. 2013. Rabies update for Latin America and the Caribbean. Emerging Infectious Diseases 19:678–679.

Walker, F. M., J. T. Foster, K. P. Drees, and C. L. Chambers. 2014. Spotted bat (*Euderma maculatum*) microsatellite discovery using illumina sequencing. Conservation Genetics Resources 6:457–459.

Walker, F. M., C. H. D. Williamson, D. E. Sanchez, C. J. Sobek, and C. L. Chambers. 2016. Species From Feces: Order-wide identification of Chiroptera from guano and other non-invasive genetic samples. Plos One 11:e0162342.

Webber, Q. M. R., R. M. Brigham, A. D. Park, E. H. Gillam, T. J. O’Shea, and C. K. R. Willis. 2016. Social network characteristics and predicted pathogen transmission in summer colonies of female big brown bats (*Eptesicus fuscus*). Behavioral Ecology and Sociobiology 70:701–712.

WHO. 2018. World Health Organization: Expert consultation on rabies: third report. World Health Organization, Geneva.

Willis, C., and R. Brigham. 2004. Roost switching, roost sharing and social cohesion: Forest-dwelling big brown bats, *Eptesicus fuscus*, conform to the fission-fusion model. Animal Behaviour 68:495–505.

Woodroffe, R., C. A. Donnelly, H. E. Jenkins, W. T. Johnston, D. R. Cox, F. J. Bourne, C. L. Cheeseman, R. J. Delahay, R. S. Clifton-Hadley, G. Gettinby, P. Gilks, R. G. Hewinson, J. P. McInerney, and W. I. Morrison. 2006. Culling and cattle controls influence tuberculosis risk for badgers. Proceedings of the National Academy of Sciences 103:14713–14717.

